# Lipids are the preferred substrate of the protist *Naegleria gruberi*, relative of a human brain pathogen

**DOI:** 10.1101/370890

**Authors:** Michiel L. Bexkens, Verena Zimorski, Maarten J. Sarink, Hans Wienk, Jos F. Brouwers, Johan F. De Jonckheere, William F. Martin, Fred R. Opperdoes, Jaap J. van Hellemond, Aloysius G.M. Tielens

## Abstract

*Naegleria gruberi* is a free-living non-pathogenic amoeboflagellate and relative of *Naegleria fowleri*, a deadly pathogen causing primary amoebic meningoencephalitis (PAM). A genomic analysis of *N. gruberi* exists, but physiological evidence for its core energy metabolism or *in vivo* growth substrates is lacking. Here we show that *N. gruberi* trophozoites need oxygen for normal functioning and growth and that they furthermore shun both glucose and amino acids as growth substrates. Trophozoite growth depends mainly upon lipid oxidation via a mitochondrial branched respiratory chain, both ends of which require oxygen as final electron acceptor. Growing *N. gruberi* trophozoites thus have a strictly aerobic energy metabolism with a marked substrate preference for the oxidation of fatty acids. Analyses of *N. fowleri* genome data and comparison with those of *N. gruberi* indicate that *N. fowleri* has the same type of metabolism. Specialization to oxygen-dependent lipid breakdown represents a hitherto unprecedented metabolic strategy in protists.

## INTRODUCTION

Members of the genus *Naegleria* are cosmopolitan protists (De Jonckheere, 2004) that dwell in fresh water. The amoebic phagotrophic stage, or trophozoite, grows by division, feeds mainly on bacteria, and occupies habitats rich in organic matter such as mud, soil, rivers, lakes and swamps (Fulton, 1970). Trophozoites can transform into a non-dividing flagellate with two flagella, but can also form dormant cysts, and can be cultured on non-bacterial food sources ranging from mammalian cell debris to non-particulate axenic culture media. By far the best known member of the genus is the thermotolerant amoeboflagellate, *Naegleria fowleri*, which causes primary amoebic meningoencephalitis (PAM) in humans, a severe and aggressive infection that usually leads to death. Cases of PAM are reported world-wide and are associated with swimming in warm waters, from which the pathogen gains access to the brain via the nasal mucosa (De Jonckheere, 2002). Trophozoites of *N. gruberi*, a nonpathogenic congener of *N. fowleri*, can be grown continuously in chemically defined media (Fulton et al., 1984). *N. gruberi* was earlier studied mainly as a model for transformation because the amoebae transform easily into flagellates, but is nowadays also used as a model to study its pathogenic relative, *N. fowleri*. Pathways of energy metabolism (core ATP synthesis) in *Naegleria* are of interest as they might harbor targets for treatment of the pathogen.

In 2010 the genome of the axenically cultured *N. gruberi* strain NEG-M was reported (Fritz-Laylin et al., 2010). The bioinformatic analyses of the genome indicated a capacity for both aerobic respiration and anaerobic metabolism with concomitant hydrogen production (Fritz-Laylin et al., 2010; Ginger et al., 2010; Opperdoes et al., 2011) and that Naegleria’s genome encodes features of an elaborate and sophisticated anaerobic metabolism (Fritz-Laylin et al., 2010). However, substrate and end product studies are still lacking. Here we investigate the ability to grow with and without oxygen of *N. gruberi* strain NEG-M and of an independent *N. gruberi* strain that was always fed with bacteria and was never grown in rich culture media. In addition, we studied the energy metabolic capacities of *N. gruberi* strain NEG-M mitochondria, its surprising spectrum of growth substrate preferences, and the pathways of ATP-production that it employs.

## RESULTS AND DISCUSSION

To test the metabolic flexibility of *N. gruberi* and — based on its genome sequence (Fritz-Laylin et al., 2010; Ginger et al., 2010; Opperdoes et al., 2011) — its anticipated capacity to switch between aerobic and anaerobic modes of metabolism, we cultured the axenic *N. gruberi* strain NEG-M under various conditions (see Methods). Under aerobic conditions the oxygen consumption of *N. gruberi* NEG-M was avid, 2.5 - 10 nmol O2 per minute per 10^6^ amoebae (Figure 1A). Trophozoites did not grow under anaerobic conditions, they became moribund and stopped multiplying in the absence of oxygen (Figure S1) or when oxygen consumption was inhibited by blocking their electron transport chain (Figure 1). Of course, cells that are cultured for years in rich culture media can lose metabolic capacities. Therefore, we also investigated a wild type strain of *N. gruberi* never grown in rich culture media (see below). This strain also stopped growing without oxygen which shows that *N. gruberi* trophozoites need oxygen for normal functioning and growth and cannot rapidly switch between aerobic and anaerobic modes of metabolism. Genome analysis led to the suggestion that *N. gruberi* could use its putative metabolic flexibility during intermittent hypoxia in their environment (Fritz-Laylin et al., 2010).

**Figure 1.**
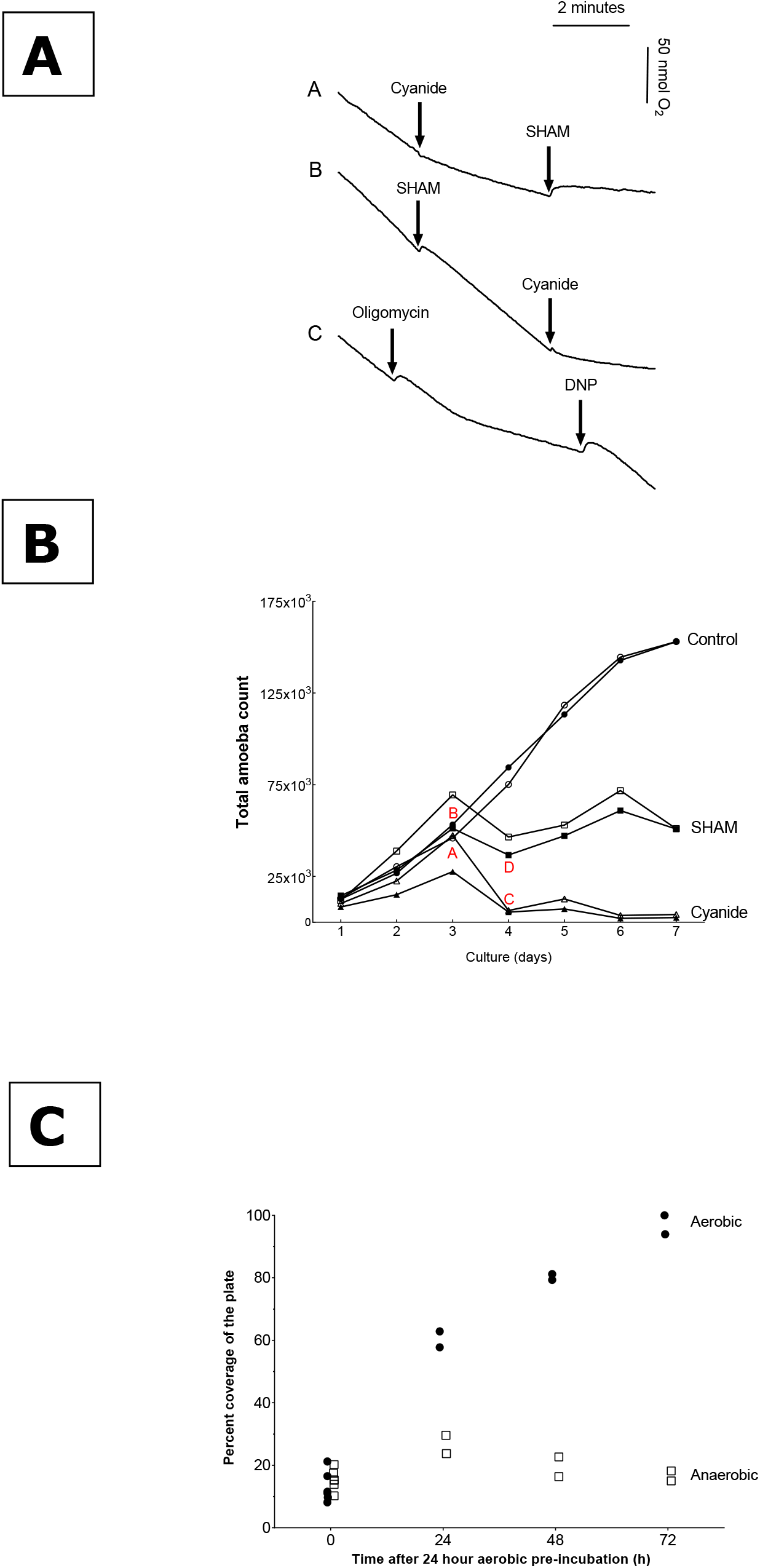
Oxygen consumption and growth under aerobic and anaerobic conditions by *N. gruberi* trophozoites. (A) Oxygen consumption of *N. gruberi* NEG-M trophozoites was measured with a Clark-type electrode. SHAM (final concentration 1.5 mM), KCN (final concentration 1 mM), oligomycine (final concentration 5 μM) and dinitrophenol (DNP, final concentration 0.1 mM) were added at the indicated time points. (B) Growth curves of *N. gruberi* NEG-M trophozoites were measured in the presence or absence of respiratory chain inhibitors. Amoebae were cultured in PYNFH for seven days. KCN (Δ, ▴) or SHAM (□, ┎) was added on day three, controls (◦, •) were treated under identical conditions. Duplicates for each growth condition are shown (open and closed symbols). Representative images of the trophozoites under the various culture conditions marked here in figure 1B with letters (A-D) are shown in Supplemental Information (See Figure S2A-D). (C) Growth of an *N. gruberi* field isolate was measured on plates covered with *E. coli* during incubations under aerobic (•) or anaerobic (□) conditions. Twelve plates were seeded with equal amounts of amoebae (as cysts) which were allowed to excyst for 24 hours under aerobic conditions at 37 °C prior to the start of the experiment. After this aerobic pre-incubation, the plates were then further incubated (starting at t=0) either aerobically or anaerobically for 24, 48 or 72 hours. The area covered by amoebae was recorded at the indicated time points.

Addition of cyanide inhibited respiration by 80% (Figure 1A, trace A), showing that mitochondrial complex IV (cytochrome *c* oxidase) was involved in aerobic growth, but also indicating that another route of oxygen consumption was operating as well. Subsequent addition of salicylhydroxamic acid (SHAM), an inhibitor of mitochondrial cyanide-insensitive plant-like alternative oxidases (AOX), resulted in a further inhibition of oxygen consumption (94%). When SHAM was added before cyanide, 14% inhibition of oxygen consumption was observed, and subsequent addition of cyanide resulted again in almost complete (94%) inhibition of the oygen consumption (Figure 1A, trace B). SHAM inhibition indicates that *N. gruberi* trophozoites use a plant-like alternative oxidase that transfers electrons from reduced ubiquinone to oxygen involving a truly branched electron transport chain (Opperdoes et al., 2011). The observed difference in degree of inhibition by cyanide and SHAM shows that complexes III and IV have a higher capacity relative to the AOX branch. Addition of cyanide or SHAM to *N. gruberi* furthermore resulted in changes in morphology and a remarkable loss of pseudopodial movement. Normal *N. gruberi* cell functions thus require both branches of the respiratory chain (see Figure 2).

**Figure 2.**
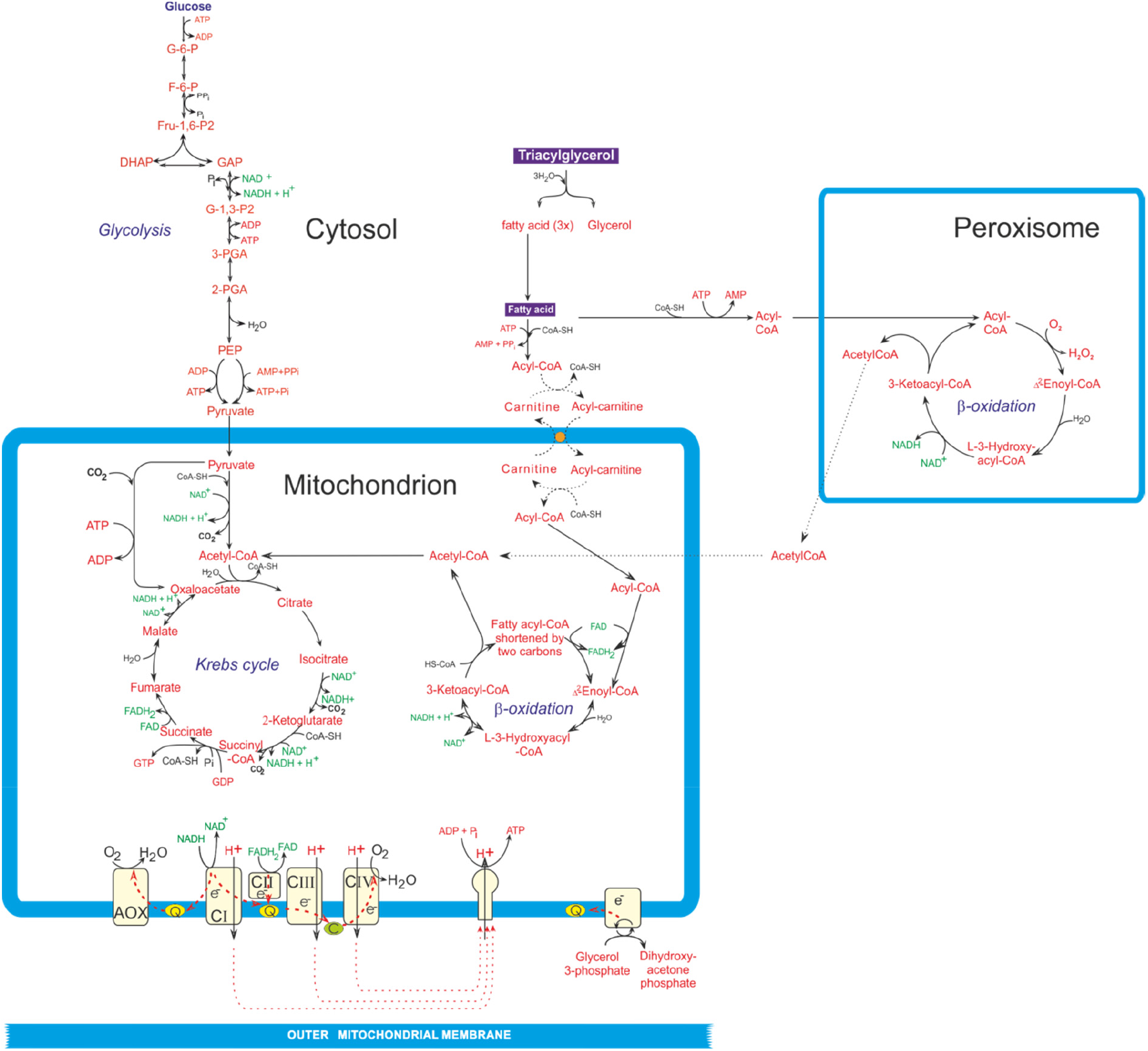
Main pathways of energy metabolism in *Naegleria gruberi* trophozoites. *Naegleria* is able to oxidize fatty acids by a process of beta-oxidation, involving both peroxisomes and mitochondria, although it is not yet clear how the chain lengths of the fatty acids are distributed over the two organelle types. It is also unknown yet how the fatty acids enter the mitochondria as up to now no carnitine acyl-CoA transferase genes have been detected in its genome. AOX, alternative oxidase; CI-IV, complexes I-IV of the respiratory chain; DHAP, dihydroxyacetone-phosphate; FAD, flavine adenine dinucleotide; FADH2, reduced FAD; F-6-P, fructose 6-phosphate; Fru-1,6-P2, fructose 1,6-bisphosphate; GAP, glyceraldehyde 3-phosphate; G-1,3-P2, glycerate 1,3-bisphosphate; G-6-P, glucose 6-phosphate; 3-PGA, glycerate 3-phosphate; 2-PGA, glycerate 2-phosphate; GTP, guanosine-triphosphate; GDP, guanosine-diphosphate; PEP, phosphoenolpyruvate. Dashed lines indicate uncertainties on the actual processes. See also Figure S3 and Tables S1 and S2.

To test cell viability under long term respiratory inhibition, we cultured *N. gruberi* NEG-M in the presence of cyanide or SHAM. The presence of cyanide resulted in a reduction of viable trophozoites by about 90% within 24 h (Figure 1B and Figures S2A and S2C), while cells treated with SHAM grew slower and merely contained fewer intracellular vacuoles (Figure 1B and Figures S2B and S2D). This shows that inhibition of the AOX is less harmful to *Naegleria* than inhibition of complex IV, which inhibits ATP synthesis, while electron overflow via AOX is necessary for *Naegleria* vitality despite having no role in ATP formation. The observed direct decrease in movement and ultimate decline of the trophozoites in the presence of cyanide, which blocks the complex III and IV branch of their electron transport chain, indicates that the mitochondrial proton gradient is the cell’s main source of ATP production. This is in agreement both with earlier observations that oxygen is essential for *N. gruberi* growth (Pittam, 1963; Weik and John, 1977b) and with our observation that oligomycin, an inhibitor of ATP synthase, strongly inhibited oxygen consumption (60%), which is again increased after addition of uncoupler (Figure 1A, trace C). Taken together, our oxygen consumption measurements and culture experiments show that *N. gruberi* trophozoites generate the bulk of their ATP via oxidative phosphorylation and that *N. gruberi* trophozoites require oxygen for homeostasis and growth, while providing no evidence for anaerobic energy metabolism.

Furthermore, we analysed the quinone content of *N. gruberi* and demonstrated that the NEG-M as well as the wild type strain contain exclusively ubiquinone and no rhodoquinone (Figure S3). Ubiquinone is used by aerobic mitochondria to transfer electrons to complex III, while rhodoquinone is used to donate electrons to a membrane-bound fumarate reductase. This reduction of fumarate to succinate is an essential reaction in malate dismutation, which is a true hallmark of the anaerobically functioning mitochondria of many eukaryotes (Tielens et al., 2002; van Hellemond et al., 1995).

To exclude the possibility that the observed strictly aerobic energy metabolism of the laboratory strain of *N. gruberi* is the result of culture-dependent metabolic losses, we also investigated the oxygen dependence of a wild type strain that was more recently isolated from river sediment and was never grown in rich culture media, but was always fed with bacteria (see Methods). For our aerobic-anaerobic growth experiment, the trophozoites of this field isolate were as ever cultured on plates covered with *Escherichia coli*, as *N. gruberi* field strains do not grow readily in liquid medium. Under aerobic conditions the plates were fully covered with amoebae after 3 days, while under anaerobic conditions no further growth was observed after the initial 24 hour growth during an aerobic incubation (Figure 1C). Thus, a field isolate of *N. gruberi*, never grown in rich culture media, is also not able to grow in the absence of oxygen. This shows that the observed obligate aerobic growth and homeostasis of the NEG-M strain does not result from loss of anaerobic capacities during decades of axenic culturing in rich media.

An oxygen-consuming respiratory chain does not necessarily imply the presence of canonical Krebs cycle activity. African trypanosomes, for example, are obligate aerobes with a branched respiratory chain comparable to that of *N. gruberi*, but despite their O_2_-dependence, procyclics of *Trypanosoma brucei* feed on glucose and produce acetate and succinate as endproducts, rather than CO_2_ and H_2_O (Besteiro et al., 2005; van Weelden et al., 2005). The existence in the *Naegleria* genome (Fritz-Laylin et al., 2010) of some enzymes considered to be typically used by eukaryotic anaerobes (Müller et al., 2012; Tielens et al., 2002), but also present in many O_2_ producing algae (Atteia et al., 2013), is interesting from an evolutionary standpoint, but only the analysis of endproducts of energy metabolism can uncover how cells actually grow.

To investigate the energy metabolism of *N. gruberi*, trophozoites were incubated in the presence of [6-^14^C]-labeled glucose, a substrate that generates labeled ^14^CO_2_ only if it is degraded via the Krebs cycle. These incubations released ^14^CO_2_ (Table 1), while no significant amounts of any other labeled endproducts could be detected by anion-exchange chromatography (not shown), demonstrating that the *N. gruberi* trophozoites degraded glucose via Krebs cycle activity without the help of fermentative pathways. The earlier genome annotations indicated already the capacity for aerobic degradation of substrates via the Krebs cycle as genes for all its enzymes are present (Fritz-Laylin et al., 2010; Ginger et al., 2010; Opperdoes et al., 2011).

**Table 1.**
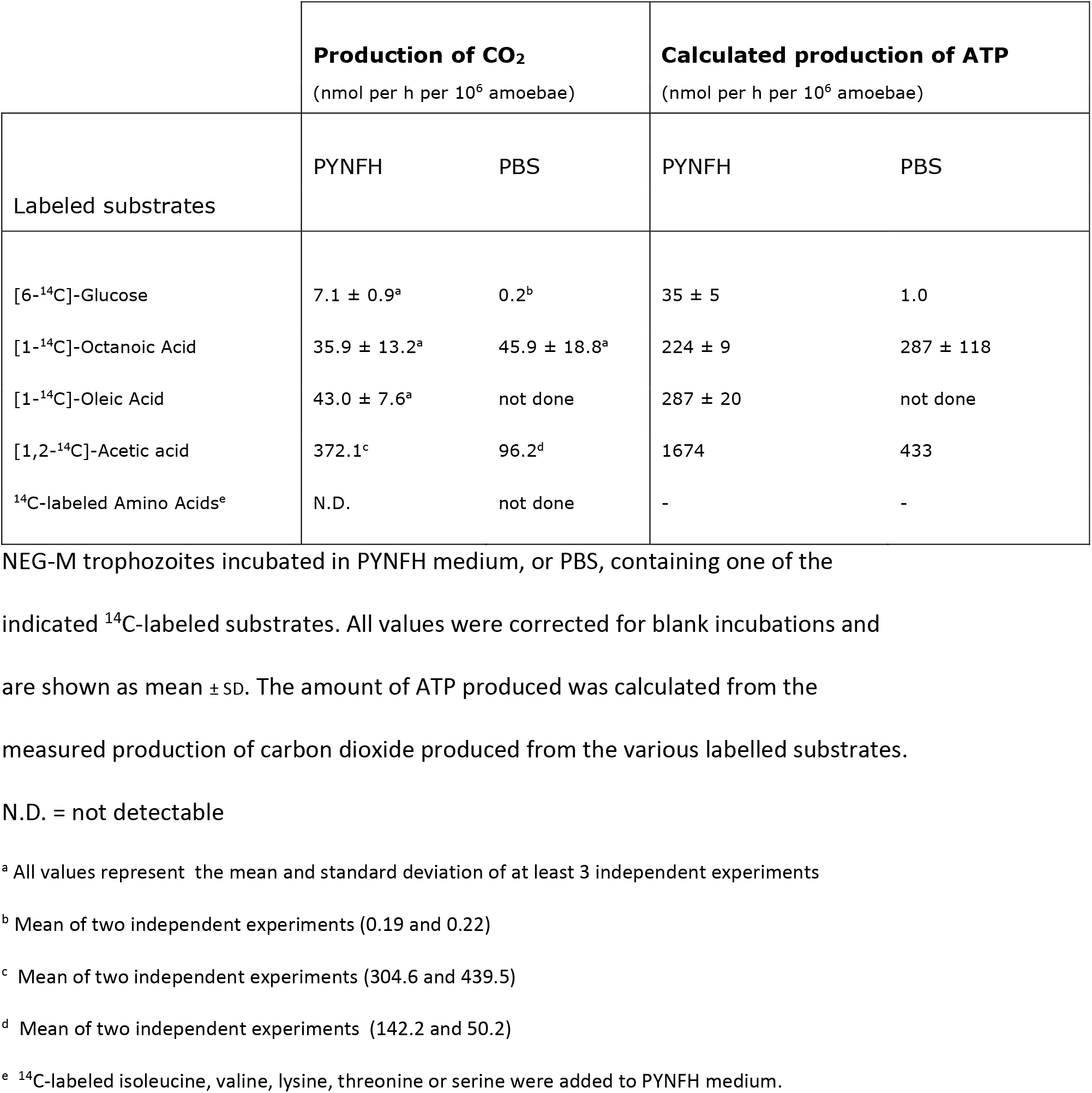
Analysis of carbon dioxide produced from ^14^C-labeled substrates by *N. gruberi*

However, the production of CO_2_ from glucose (7 nmol per hour per 10^6^ amoebae) accounted for only 1-5% of the overall O_2_ consumption, which was around 150-600 nmol per hour per 10^6^ amoebae (Figure 1A). Clearly, oxygen was mainly consumed by oxidation of substrates other than glucose present in the very rich PYNFH culture medium because in standard aerobic degradation of glucose to carbon dioxide, O_2_ consumption and CO_2_ production occur in 1:1 stoichiometry (glucose + 6 O_2_ ➔ 6 CO_2_ + 6 H_2_O).

To determine whether removal of alternative substrates could stimulate glucose consumption, we incubated the trophozoites in phosphate-buffered saline with 5 mM glucose as the sole carbon source. Under these conditions, CO_2_ production was even lower than in the rich PYNFH medium (Table 1).

To see if *N. gruberi* degrades proteins present in PYNFH medium for amino acid oxidation, the full set of enzymes for which is present in the *N. gruberi* genome (Opperdoes et al., 2011), we incubated trophozoites in PYNFH medium supplemented with one of the following ^14^C-labeled amino acids: isoleucine, valine, lysine, threonine or serine. After 24 hours, no ^14^CO_2_ release at all could be detected in any of these incubations (Table 1). Thus amino acids are also not the amoeba’s growth substrate.

Incubations in the presence of ^14^C-labeled oleic acid (C_18:1_) or octanoic acid (C_8:0_) resulted in the production of ^14^CO_2_ in amounts five- to six-fold higher than during the incubations with labeled glucose (Table 1). Analysis of the incubation medium by anion exchange chromatography revealed no significant amounts of metabolic endproducts other than CO_2_. We also tested ^14^C-labeled acetate as a direct precursor for Krebs cycle activity. Acetate was rapidly degraded to ^14^CO_2_ (Table 1), which confirms the activity of the Krebs cycle.

The combined data indicate that *Naegleria* trophozoites prefer fatty acids to both glucose and amino acids as growth substrates. They oxidize fatty acids and degrade the resulting acetyl-CoA units via the Krebs cycle. In many eukaryotes, carbohydrates and most amino acids can be degraded by anaerobic fermentations to endproducts such as lactate, acetate, propionate and succinate (Müller et al., 2012). Fatty acids are, however, non-fermentable substrates and their oxidation requires the mitochondrial respiratory chain.

To more thoroughly exclude the possible use of other pathways and to independently confirm, via a different analytical technique, the use of fatty acids by *N. gruberi*, we incubated the NEG-M strain in the presence of ^13^C-octanoic acid and analysed the resulting incubation medium by NMR spectroscopy (Figure 3). Comparison of the signal intensity of the peaks of [^13^C]-labeled carbon atoms 2, 3 and 4 of octanoic acid of the incubation that contained *N. gruberi* NEG-M amoebae (top graph) with the control incubation (bottom graph) showed that in the incubations containing trophozoites, octanoic acid was readily consumed (60% was degraded), while no ^13^C-labeled fermentation products such as acetate, propionate or succinate could be detected (Müller et al., 2012), which is in full agreement with our ^14^C-labeled fatty acid incubations.

**Figure 3.**
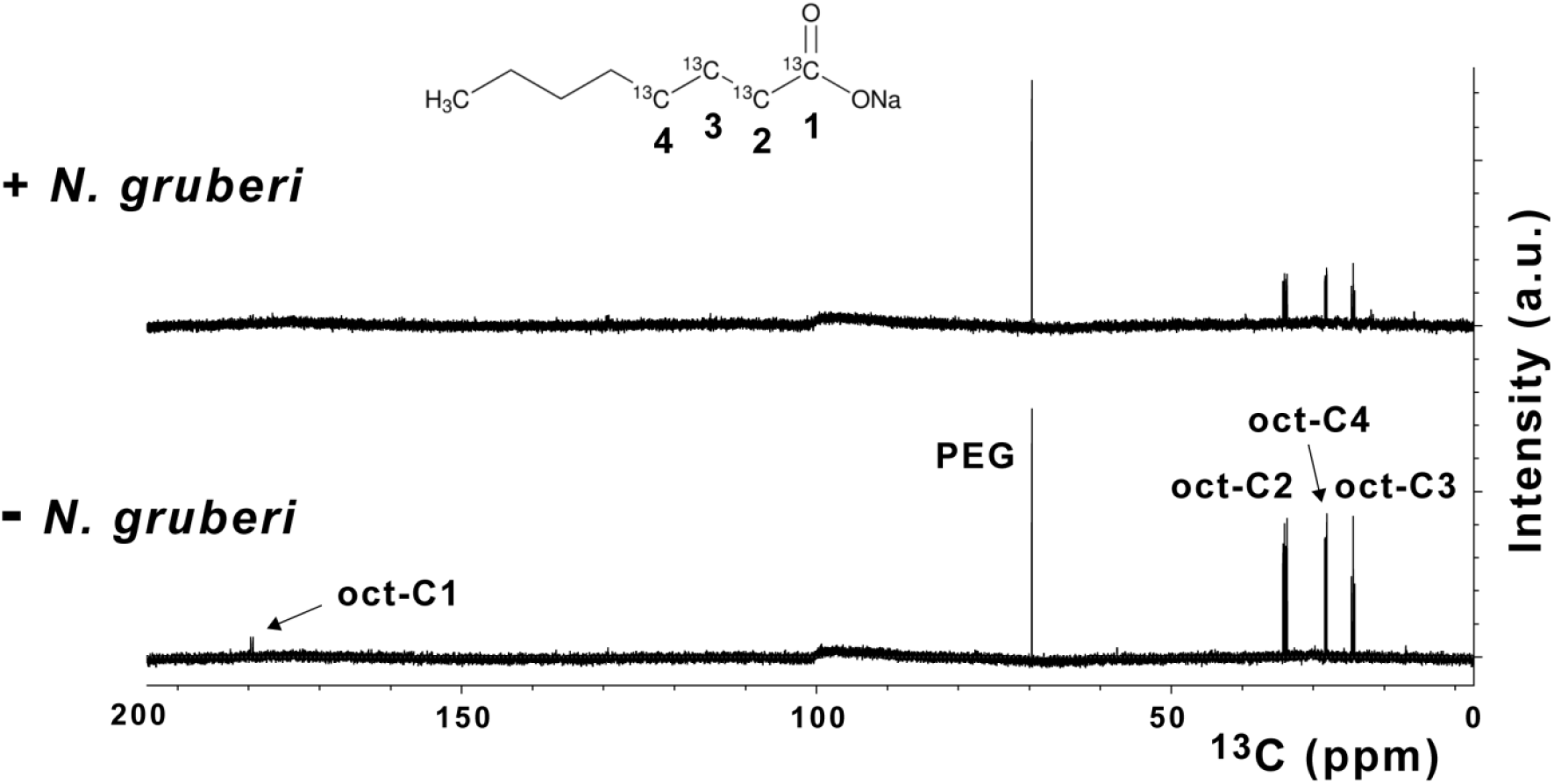
NMR analysis of [^13^C]-labeled octanoic acid degradation by *N. gruberi*. Trophozoites were incubated in phosphate buffered saline (PBS) containing 45 μM polyethylene glycol 6000 (PEG) plus 210 μM [1,2,3,4-^13^C]-labeled octanoic acid. Identity of the peaks originating from the four labeled carbon atoms of octanoic acid is indicated by C1-C4.

Parasites are known to live on fermentation of sugars, whether they live in blood, organs or the gut (Köhler and Tielens, 2008). Eukaryotic pathogens that invade human tissues but do not live primarily on glucose, have not been reported to date. *Naegleria* presents an unprecedented example of a human pathogenic protist that shuns glucose as a carbon and energy substrate. However, our observation that when given a choice, lipids are the preferred substrate, does not imply that other substrates, such as glucose, are not used to a lower extent at the same time, which is in fact what we observed (Table 1). Furthermore, the observation that lipids are the preferred substrate, does not imply that *N. gruberi* cannot grow in its absence.

The present findings allow us to propose a metabolic map for the energy metabolism of *N. gruberi* trophozoites that reflects the obligate aerobic nature of the growing organism and its reliance on lipid oxidation for energy metabolism (Figure 2). Though *N. gruberi* contains all the genes required for the metabolism of carbohydrates and amino acids (Opperdoes et al., 2011), when given the choice during growth it exhibits a clear preference for fatty acid oxidation as the main source for ATP production (Table 1). The genome encodes a full set of lipid-degrading enzymes, including lipases, phospholipases and β-oxidation enzymes (Opperdoes et al., 2011) (Table S1) that underpin Figure 2. A close inspection of sequences of proteins involved in lipid degradation and the beta-oxidation of the liberated free fatty acids that are encoded in the *N. gruberi* genome revealed that the genome contains many sequences with peroxisomal targeting signals (Opperdoes et al., 2011) in addition to isofunctional homologues that carry a predicted mitochondrial transit sequence (Table S1). This suggests that both peroxisomes and mitochondria are involved in the oxidation of fatty acids.

Fatty acids can not however be the exclusive substrates for *Naegleria* growth, as degradation of acetyl-CoA to carbon dioxide requires the presence and occasional replenishment of Krebs cycle intermediates and these cannot be made from fatty acids. This implies that complete oxidation of fatty acids to carbon dioxide requires a trickle of carbohydrate oxidation to pyruvate, which fits in the observed metabolism of glucose and fatty acids by *N. gruberi* (Table 1). The preferential use of lipids instead of carbohydrates is also reflected in the storage of substrates. Early histochemical studies of *N. gruberi* showed that spherical globules (0.4-1.0 μm) consisting of lipids are distributed throughout the cytoplasm, whereas glycogen is absent (Pittam, 1963). Parasites are known to obtain lipids from their host for usage as building blocks and increased expression of enzymes involved in lipid metabolism under certain conditions or in various life-cycle stages has also been observed in transcriptome and proteome studies (Atwood et al., 2005; Li et al., 2016; Roberts et al., 2003; Saunders et al., 2014; Trindade et al., 2016; Yichoy et al., 2011). Substrate and endproduct studies are lacking, however, and the preferential use of lipids as the major bioenergetic substrate has not been shown for any eukaryotic pathogen to date. In nature *N. gruberi*, like many other protists, feeds mainly on bacteria, a food source containing larger amounts of lipids than carbohydrates. Other phagotrophic protists might be found that exhibit a similar substrate preference for lipids. Furthermore, parasitic protists often encounter large variations in their menu during the life cycle and the metabolism of many parasitic protists is therefore very flexible.

Based on our analysis of the genome content of *N. fowleri* and comparison with that of *N. gruberi* (Table S2), we suggest that the strong preference for lipids should also be found in the thermophilic congener, *N. fowleri*, which exhibits a strong tropism for the olfactory nerves of the nasal mucosa that it infects en route to the brain during PAM. The white matter of brain is rich in myelin sheets, which surround the axons of nerve cells and contain about 80% lipids by weight (O’Brien and Sampson, 1965). *N. fowleri* encodes a set of glycohydrolases, glycoside hydrolases, glucosyl ceramidase, sphingomyeline phosphodiesterase and phospholipases, several present in multiple copies, that could channel myelin sheaths into its energy metabolism (Table S2). Acetate, here shown to be readily consumed by *Naegleria*, is also amply available in brain (Wyss et al., 2011). Though the preferred growth substrates of the “brain-eating amoeba” as the deadly and more difficult to culture pathogen *N. fowleri* is sometimes called, have still not been directly shown, an earlier investigation suggested that they possibly are not carbohydrates (Weik and John, 1977a). The endproducts of glucose and fatty acid degradation reported here provide a clear picture for *N. gruberi* and make very specific predictions for *N. fowleri* in agreement with Weik and John’s (1977a) suggestion. Fatty acid oxidation in *Naegleria* uncovers a novel energy metabolic specialization in protists with evolutionary and medical significance and could point the way to improved treatments for PAM.

## EXPERIMENTAL PROCEDURES

### *Naegleria* strains

*Naegleria gruberi* strain NEG-M (ATCC^®^ 30224™) was cultured at 25 °C while shaking at 50 rpm in PYNFH medium, which contains Peptone, Yeast extract, Nucleic acid, Folic acid, Hemin and 10% heat-inactivated fetal bovine serum (ATCC medium 1034). This medium was supplemented with 40 μg/ml gentamicin, 100 units/ml penicillin and 100 μg/ml streptomycin. A field isolate of *N. gruberi* was obtained from sediment in Rio Verde, Tuzigoot, Arizona, USA (De Jonckheere, 2007). This field isolate has the same ITS rDNA sequence as strain NEG-M, indicating that it is the same species (De Jonckheere, 2014). Since its isolation from the field this *N. gruberi* isolate (FI trophozoites) was always cultured on nonnutrient agar plates coated with *Escherichia coli* and this strain was never cultured in any of the culture media that are used to grow the NEG-M strain. For our experiments the amoebae were cultured at 37 °C on non-nutrient agar plates seeded with Top10 *E. coli* (ThermoFisher Scientific). Amoebae were counted using Differential Interference Contrast (DIC) microscopy and images taken were analyzed using Cell^^^F Software (Olympus).

### Trophozoite culture of *N. gruberi* NEG-M in the absence of oxygen or in the presence of inhibitors of respiration

To investigate the anaerobic capacity of *N. gruberi* NEG-M, trophozoites were seeded at a density of 30,000 trophozoites per ml in 5 ml PYNFH medium containing 40mM ascorbate (anaerobic incubations) or 40mM NaCl (aerobic incubations). In aerobic incubations the gas phase was air. For anaerobic incubations the medium was degassed, flasks were flushed for 5 minutes with nitrogen and prior to closing the bottles, ascorbate oxidase (100 U) was added to the anaerobic incubations to remove any remaining oxygen (Tielens et al., 1981).

To determine the effects of respiratory chain inhibitors on the cell growth of *N. gruberi* NEG-M, trophozoites were seeded in PYNFH medium at a density of 30,000 trophozoites per ml. Upon reaching log growth (day 3) respiratory chain inhibitors were added, KCN was added to the culture in a final concentration of 1 mM, SHAM was dissolved in 96% ethanol and added to the culture at a final concentration of 1.5 mM SHAM, 0.5% ethanol. Control incubations contained 0.5% ethanol. Throughout cell growth experiments, the medium was changed daily, and KCN and SHAM were added fresh daily.

### Anaerobic trophozoite culture of *N. gruberi* field isolate

Fresh NN-plates were inoculated by placing a circular slice (9 mm in diameter) of an agar plate covered uniformly with *N. gruberi* cysts upside down on the center of the plate. These inoculated plates were then pre-incubated aerobically for 24 hours at 37°C to allow excystment of amoebae. After this aerobic pre-incubation the amoebic growth perimeter was marked and recorded, and the plates were then further incubated aerobically or anaerobically for 72 hours. Anaerobic conditions were created by placing the plates in gas-tight jars in which the air was replaced by a mixture of 85% N2, 10% CO_2_, 5% H2 using an Anoxomat (Mart Microbiology, Drachten, The Netherlands), aerobic plates were incubated under normal atmospheric conditions in identical jars. After 24, 48 and 72 hours an anaerobic and an aerobic jar, each containing two plates, were opened and the perimeter of growth on the anaerobic and aerobic plates was recorded.

### Respiration of N. *gruberi* NEG-M

Oxygen consumption by *N. gruberi* NEG-M trophozoites was determined using a Clark-type electrode at 25 °C in 2 ml oxygen-saturated fresh medium. KCN and SHAM were used at a final concentration of 1 mM and 1.5 mM, respectively.

### Quinone-composition of *N. gruberi* NEG-M

*N. gruberi* NEG-M was cultured in PYNFH medium and harvested during logarithmic growth. Lipids were isolated by a chloroform/methanol extraction procedure (Bligh and Dyer, 1959). Ubiquinone and rhodoquinone content was analysed by a Liquid Chromatography-Mass Spectrometry (LC-MS) method as described before (van Hellemond et al., 1995). Quinones from *Fasciola hepatica* were isolated using the same method and analysed as a positive control. The quinone composition of the *N. gruberi* field isolate was analysed using the same procedure.

### Metabolism of *N. gruberi* NEG-M

Trophozoites were harvested during logarithmic growth and 3.5×10^6^ trophozoites were transferred to a sealed 25 ml erlenmeyer flask containing either 5 ml PYNFH medium or phosphate buffered saline (PBS). The incubation was started by addition of one of the labeled substrates (all supplied by PerkinElmer, Boston, MA, USA): D-[6-^14^C] glucose (5 mM, 5 μCi), [1-^14^C] octanoic acid (210 μM, 5 μCi), [1-^14^C] oleic acid (210 μM, 5 μCi), [1,2-^14^C] acetic acid (5 mM, 5 μCi), [U-^14^C] isoleucine (5 μCi), [U-^14^C] valine (5 μCi), [U-^14^C] lysine (5 μCi), [U-^14^C] threonine (5 μCi) and [3-^14^C] serine (5 μCi). Blank incubations without trophozoites were started simultaneously. All samples were incubated at 22 °C while shaking gently at 125 rpm. After 18-24 hours the incubations were terminated and endproducts were analysed as described before(van Weelden et al., 2003). In short, 100 μl NaHCO_3_ (25 mM) was added and the incubation was ended by addition of 6 M HCl, lowering the pH to 2.5. Carbon dioxide was trapped in a series of four scintillation vials, each filled with 1 ml of 0.3 M NaOH and 15 ml of Tritisol scintillation fluid(Pande, 1976). Thereafter the radioactivity in this fraction was counted in a scintillation counter. After this removal of carbon dioxide, the acidified supernatant was separated from the cells by centrifugation (4 °C for 10 min at 500×g) and neutralized by the addition of NaOH. The labelled end-products were analysed by anion-exchange chromatography (Tielens et al., 1981). Fractions were collected and radioactivity was measured in a scintillation counter after addition of LUMA-gel (Lumac*LCS, Groningen, The Netherlands).

NMR spectroscopy was used to analyse samples that had been incubated in phosphate buffered saline (PBS) containing 45 μM polyethylene glycol 6000 (PEG) plus 210 μM ^13^C-labeled [1,2,3,4-^13^C]-labeled octanoic acid (Cambridge Isotope Laboratories, Andover, MA, USA) for 24 h at 22 °C while shaking gently at 125 rpm. After the incubations the cells were spun down (4 °C for 10 min at 500×g) and 500 μl of the supernatants were analysed. NMR experiments were performed at 25 °C on a 600 MHz Bruker Avance spectrometer equipped with cryogenically cooled TCI-probe for extra sensitivity. ^13^C experiments were recorded from 0-200 ppm employing power-gated proton decoupling. 16k FIDs were accumulated with interscan delay of 2 seconds, and acquisition times of 0.5 s (32k complex points). Processing was performed with exponential multiplication and Fourier Transformation, followed by base-line correction.

## Supplemental Information

Supplemental Information includes three figures and two tables.

## Acknowledgements

We thank Matthias Wittwer (Spiez Laboratory, Federal Office for Civil Protection, Spiez, Switzerland) for making available the annotated transcriptome sequences of *Naegleria fowleri* ATCC 30863 trophozoites. W.F.M. thanks the ERC (grant 666053) for funding and H.W. acknowledges financial support for the NMR experiments by NWO-Groot Grant 175.107.301.10.

## Author Contributions

M.L.B., J.J.vH. and A.G.M.T. conceived and designed the experiments. M.L.B. and M.J.S. conducted the experiments. F.R.O., V.Z. and W.F.M performed comparative genomic analyses. J.F.DJ. provided the field isolate of *N. gruberi* and know how on amoebae. J.J.vH. and J.F.B. developed and performed the quinone analyses. H.W. performed the NMR analysis. J.J.vH and A.G.M.T. supervised the project. M.L.B., F.R.O., W.F.M. and A.G.M.T. wrote the manuscript. All authors participated in discussion, commenting on and final approval of the manuscript.

